# Little Evidence for Homoeologous Gene Conversion and Homoeologous Exchange Events in *Gossypium* Allopolyploids

**DOI:** 10.1101/2023.11.08.566278

**Authors:** Justin L Conover, Corrinne E Grover, Joel Sharbrough, Daniel B Sloan, Daniel G Peterson, Jonathan F Wendel

## Abstract

A complicating factor in analyzing allopolyploid genomes is the possibility of physical interactions between homoeologous chromosomes during meiosis, resulting in either crossover (homoeologous exchanges) or non-crossover products (homoeologous gene conversion). This latter process was first described in cotton by comparing SNP patterns in sequences from two diploid progenitors with those from the allopolyploid subgenomes. These analyses, however, did not explicitly account for autapomorphic SNPs that may lead to similar patterns as homoeologous gene conversion, creating uncertainties about the reality of the inferred gene conversion events. Here, we use an expanded phylogenetic sampling of high-quality genome assemblies from seven allopolyploid *Gossypium* species (all derived from the same polyploidy event), four diploid species (two closely related to each subgenome), and a diploid outgroup to derive a robust method for identifying potential genomic regions of gene conversion and homoeologous exchange. Using this new method, we find little evidence for homoeologous gene conversion in allopolyploid cottons and that only two of the forty best-supported events are shared by more than one species. We do, however, reveal a single, shared homoeologous exchange event at one end of chromosome 1, which occurred shortly after allopolyploidization but prior to divergence of the descendant species. Overall, our analyses demonstrate that homoeologous gene conversion and homoeologous exchanges are uncommon in *Gossypium*, affecting between zero and 24 genes per subgenome (0.0 - 0.065%) across the seven species. More generally, we highlight the potential problems of using simple four-taxon tests to investigate patterns of homoeologous gene conversion in established allopolyploids.

**SIGNIFICANCE STATEMENT:** Allopolyploidy is a prominent process in plant diversification, involving the union of two divergent genomes in a single nucleus via interspecific hybridization and genome doubling. The merger of genomes sets in motion a variety of inter-genomic and epigenomic interactions that are thought to lead to the origin of new phenotypes. Among these is recombinational exchange between duplicated chromosomes, which can involve sequence lengths ranging from several bases to entire chromosome arms, and which can be either reciprocal or unidirectional in their effects. Here we present a new analytical framework for detecting these inter-genomic recombinational processes in allopolyploids, and demonstrate that they have been rare in a group of allopolyploid species in the cotton genus.

## INTRODUCTION

Whole genome duplication events (polyploidy) are a prominent force in the evolution of plants. Polyploidy is exceptionally common in angiosperms, where it is an active ongoing process in many lineages. In addition, all angiosperms have a deep phylogenetic history that includes on average three or four rounds of polyploidy events (1), including one event that is shared by all flowering plants (2). Polyploidy has also played an important role in crop species, as many economically important crops are either currently polyploid (*e.g*., cotton, quinoa, potatoes) or have experienced a polyploidization event in their recent evolutionary pasts (*e.g*., maize, *Brassica* crops (3, 4)). Understanding the dynamics of genome evolution following polyploid formation is therefore important to our understanding of plant evolution, but also has important potential economic and agricultural consequences.

One of the complicating factors in studying the genomes of allopolyploids (*i.e*., polyploids that arise through merger of divergent genomes (5, 6)) is the possibility for physical interactions between their two (or more) co-resident genomes (*i.e*., subgenomes) during meiosis. During the process of double-strand DNA break repair, the broken strand of one chromatid can be repaired using its homoeologous (rather than homologous) chromosome copy. If this repair includes chiasma formation between homoeologous chromosomes (indicated by the formation of multivalents), then recombination is expected to lead to homoeologous exchanges (HEs) that reciprocally affect the region of the chromosome arm located between the chiasma and telomere (reviewed in (7)). The resulting haplotype blocks of HEs are then broken up in subsequent generations via homologous recombination (and/or lost via drift), making the detection of these regions generally difficult and potentially affecting each subgenome unequally. Double-stranded breaks may also be repaired via non-crossover pathways involving the homoeologous chromosome, resulting in homoeologous gene conversion (hGC, also known as nonreciprocal homoeologous recombination (8)), which distort Mendelian segregation patterns (9). Blocks of hGC are localized to the initial site of the double-stranded break and, although little is known about the typical length of hGC blocks, are substantially shorter than those resulting from HE events. Studies of homologous gene conversion in diploid *Arabidopsis* suggests that typical gene conversion blocks can range in size from tens to thousands of base pairs in length and do not appear to be biased towards creating higher GC content (10), contrary to patterns of homologous gene conversion in other eukaryotes (9, 11).

Ultimately, both HE and hGC can act to homogenize sequences across otherwise divergent subgenomes, thereby complicating allopolyploid genome assembly and analyses. In turn, this sequence homogenization can generate heterogeneous phenotypes (as demonstrated in *Brassica* (12, 13), *Tragopogon* (14, 15), *Oryza* (16–18)) by altering allele dosage and other (epi)genomic patterns (19), acting to reshuffle genetic variation and potentially resulting in novel genomic combinations for selection to act upon. Although multiple methods have been implemented to identify regions of allopolyploid genomes that have experienced homoeologous exchanges (e.g. competitive read mapping (20), ABBA-BABA tests (21), phylogenomics (22), or chromosome staining (15)), comparatively little attention has been paid to developing methods that can identify homoeologous gene conversion events.

Homoeologous gene conversion was first described in allopolyploid cotton (8) using expressed sequence tags (ESTs) and employing an analytical method similar to those developed to identify gene conversion in highly heterozygous diploids (most commonly created by crossing divergent, highly inbred lines) (10). In short, EST alignments were generated to include an allopolyploid cotton species (represented by both subgenomes) and its two model diploid progenitors, wherefrom homoeoSNPs (*i.e*., SNPs that distinguish one subgenome and its closest diploid progenitor from the other subgenome and its diploid progenitor) were identified and treated as analogous to the SNPs traditionally used in diploid-based investigations. Using this method, Salmon et al., (8) found that hGC may affect as many as 1-2% of genes in cotton. This estimate was later updated (23) to include additional members of the polyploid clade and using a more extensive EST dataset, finding that approximately 7% of genes have been affected in one or both allopolyploid species evaluated. Subsequent efforts in evaluating hGC in cotton have relied on similar logic, albeit extending this to the increasingly available high-throughput sequences (24–27) including full genomes, all of which suggest gene conversion in allopolyploid cottons is relatively rare.

Despite the clear rationale of extending these diploid individual-based ‘quartet’ methods to an evolutionary perspective in a polyploid context, there remain a number of potential problems noted with this approach (see *e.g*., (8, 23, 27)) that have not been explored. The most difficult to address is that SNP patterns indicative of hGC or HE could also arise via other evolutionary mechanisms. For example, the canonical 3:1 SNP pattern of quartet-based analyses could also be caused by (1) mutations that occur in one diploid lineage after its divergence from the actual parental lineage to the allopolyploid, or (2) mutations that occur in the common ancestor of one diploid/subgenome lineage followed by another mutation at that same site in the other subgenome of the polyploid. As such, differentiating between the various evolutionary processes that can give rise to hGC/HE-like SNP patterns is important but nearly impossible without estimating the rates at which these autapomorphic and homoplasious SNP patterns occur. Further exacerbating this problem, HE/hGC SNP patterns are expected to become more common in older polyploids as the evolutionary distance between the polyploid subgenomes and their diploid relatives diverge. It is important to note that the evolutionary distance between diploids and their derivative allopolyploid subgenomes need not be equivalent, and thus the rates of autapomorphic and homoplasious SNPs need not be equal between the two lineages of the allopolyploid subgenomes. For example, if the closest extant diploid relatives to an allopolyploid differ in their relatedness to their polyploid subgenomic counterparts (*i.e.*, if one ‘true’ diploid progenitor goes extinct shortly following allopolyploid formation), then the terminal branch of the phylogenetic tree leading to the extant diploids would differ, and more autapomorphic SNPs would be expected to occur on the longer terminal branch. Therefore, refining the methodology to differentiate between the number of homoeoSNPs that are truly caused by hGC or HE from those caused by autapomorphic or homoplasious SNPs, as well allowing for different rates of autapomorphic or homoplasious SNPs between the two lineages, is an important area of improvement for the analysis of HE and hGC.

*Gossypium* is an ideal system (28–30) to develop analytical methods for detecting hGC and HE events in allopolyploids. The genus contains ∼45 currently recognized diploid species, which are classically categorized into eight genome groups (named A-G, K) based on genome size, karyotype, and patterns of intercompatibility (30–32). The genus also includes seven allopolyploids (named the AD clade), all of which are descended from the same polyploidization event (33), which occurred via hybridization 1-2 million years ago between a member of the D-lineage (most closely related to *G. raimondii* (D5)) and a member of the A genome group (equally related to *G. arboreum* (A2) and *G. herbaceum* (A1)). Economic interest in the cotton genus led to the development of high-quality genome resources for multiple species, with chromosome-scale genomes available for all seven (34, 35) allopolyploids (including multiple sequences of the domesticated *G. hirsutum* and *G. barbadense* (*36*)), 10 diploids representing the diversity of the genus, and an outgroup to the genus, *Gossypioides kirkii*. Included within the diploid genome assemblies are the extant model diploid progenitors and their closely related diploid outgroups (37, 38), allowing for powerful analyses of post-polyploidization genome evolution. Finally, there is little chromosome number evolution within *Gossypium*, with all diploid species containing 13 chromosomes (*Gossypioides kirkii* has n=12), thereby simplifying the process of developing whole-genome alignments, even in the face of a two-fold difference in genome size between the diploid lineages that gave rise to the allopolyploid. Finally, there are hitherto no reported regions of homoeologous exchange in any allotetraploid species, the presence of which could make differentiating hGCs from HEs difficult.

Here, we leverage multiple high-quality genome sequences within *Gossypium* to evaluate the extent to which we can disentangle hGC and HE from other evolutionary phenomena capable of producing similar patterns, extending the previous analyses of hGC in *Gossypium* to all seven monophyletic allopolyploid species. Using a well-established phylogenetic framework, we develop a robust methodology to identify potential hGCs/HE events, finding little evidence that hGC occurs in any *Gossypium* allopolyploid lineage. Nevertheless, we describe a small number of regions (∼40 in total across all seven polyploids) that may have experienced hGC, HEs, or other mechanisms of inter-subgenomic sequence translocation, including the first described instance of homoeologous exchange in *Gossypium*. We discuss the implications of our work for other analyses of hGC and highlight the misleading results that may be obtained using the ‘quartet’ method to identify hGC, especially in older allopolyploids where autapomorphic SNPs are likely and in situations where the extant diploids are not closely related to the allopolyploid subgenomes. We also discuss the role that genome assembly quality plays in the ability to identify potential hGC events.

## RESULTS

### An Improved Method to Identify Potential Homoeologous Gene Conversion (hGC) Events

Homoeologous gene conversion was initially described in allopolyploid cotton (as non-reciprocal homoeologous recombination, or NRHR) by comparing alignments of EST sequences from the two model diploid progenitor diploid species (*G. raimondii* and *G. arboreum*) with orthologous EST sequences from the two subgenomes of allopolyploid *G. hirsutum* (Figure 1A). For the sake of generalization, we arbitrarily designate *G. raimondii* as D_1_ (*i.e*., diploid species 1), *G. arboreum* as D_2_ (*i.e*., diploid species 2), and the two subgenomes of an allotetraploid as P_1_ (the subgenome most closely related the D_1_) and P_2_ (the subgenome most closely related to D_2_). HomoeoSNPs were first identified as those positions where the diploid progenitors contained nucleotides each matching their respective subgenome (*i.e*., D_1_=P_1_ and D_2_=P_2_). Subsequently, diagnostic SNP patterns that may indicate HE or hGC were identified in those sites where the polyploid subgenomes were both equivalent to each other and different from one of the two parental diploids. Specifically, sites where the D_1_ diploid contained a different nucleotide from that shared by D_2_ and both the P_1_ and P_2_ subgenomes (*i.e*., D_2_=P_2_=P_1_) were considered putative HE/hGC sites in which the P_2_ subgenome had “overwritten” the P_1_ subgenome (Figure 1A, red box). Likewise, sites in which D_2_ contained one allele, while D_1_, P_1_, and P_2_ shared a second allele (*i.e*., D_1_=P_1_=P_2_), were considered putativeHE/hGC events where the P_1_ subgenome had overwritten the P_2_ subgenome (Figure 1A, blue box). Finally, we note that the logic describing the diagnostic SNP patterns is equivalent for hGCs as it is for HEs. Therefore, for the sake of simplicity, we will only refer to hGC SNP patterns throughout the results (except for the section below describing a shared HE event) and explore the difficulties in differentiating hGC from HE in the discussion.

**Figure 1:**
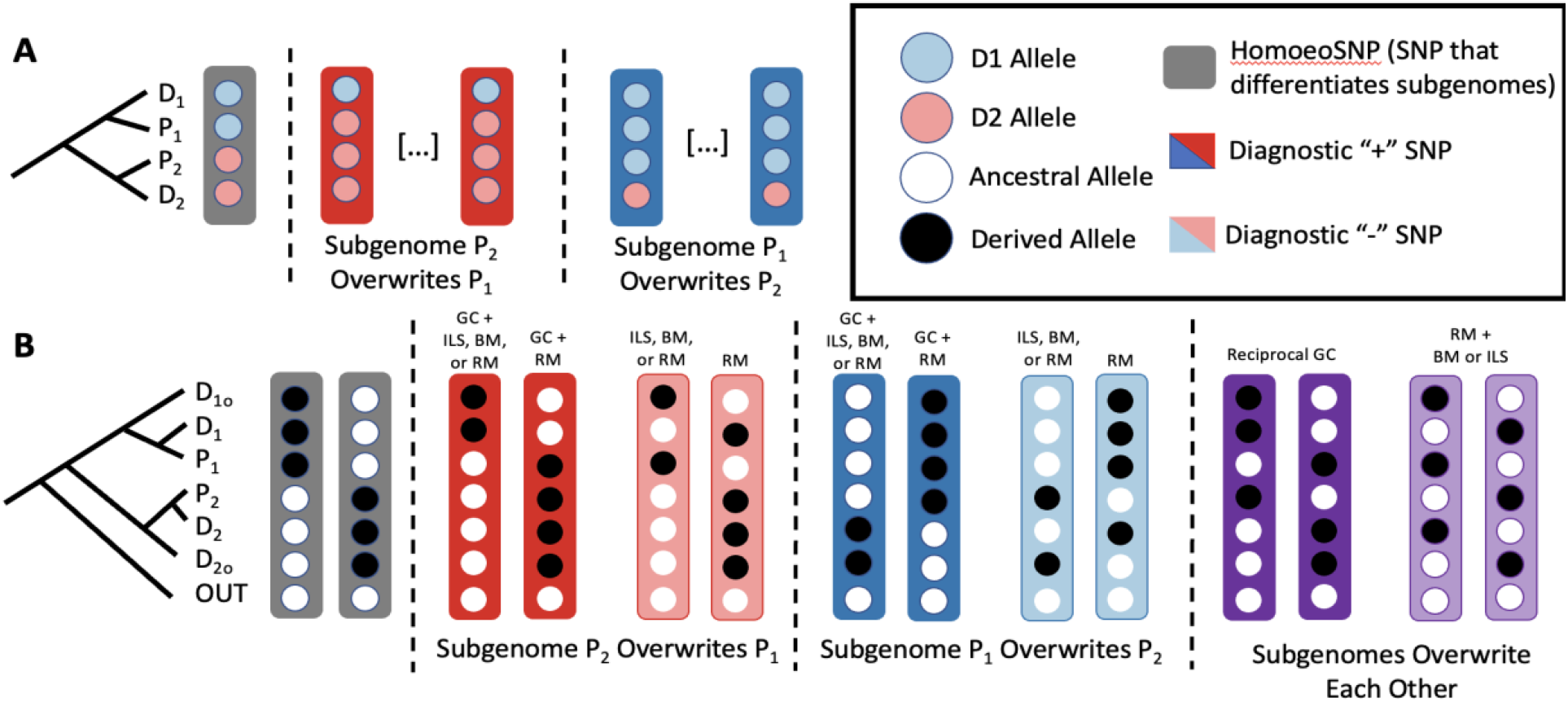
Overview of Methods to Identify Gene Conversion Events. **(A)** Classical method of identifying hGC events. Given a four taxon tree consisting of two model diploid progenitors (D_1_ and D_2_) and the two subgenomes of an allopolyploid (P_1_ and P_2_), categories of SNP patterns have been used to infer patterns of hGC. Blue circles indicate SNPs that are found in the D_1_ diploid, and red circles indicate SNPs found in the D2 diploid. First, SNPs that can reliably differentiate one diploid/subgenome clade from the other (homoeoSNPs; gray box) are identified. Between consecutive homoeoSNPs, if multiple SNPs are consistent with the pattern where subgenome P1 has overwritten P2 (blue box), or vice versa (red box), then gene conversion is inferred. Importantly, however, there are other evolutionary forces that may generate these SNPs patterns (*e.g*., autapomorphic SNPs in either diploid terminal branch) that are not accounted for in this model. **(B)** Our model for investigating rates of homoeologous gene conversion and homoeologous exchanges. Given a seven taxon tree consisting of an outgroup (OUT, used to polarize SNPs as ancestral or derived), subgenomes of an allotetraploid (P_1_ and P_2_, respectively), two diploid progenitor species (D_1_ and D_2_, respectively), and an outgroup to each of the diploid/subgenome clades (D_1o_ and D_2o_, respectively), we define homoeoSNPs as those SNP sites where either the (D_1_, D_1o_, P_1_) or the (D_2_, D_2o_, P_2_) clade exclusively have derived SNPs. SNP sites that are potentially the result of hGC (Diagnostic + SNPs) are highlighted in dark blue (P_1_ overwrites P_2_) or dark red (P_2_ overwrites P_1_). For each of these SNP patterns, there are multiple other mutational/evolutionary patterns that may result in the same pattern, including recurrent mutation (*i.e*., mutation on internal branch leading to D_1_/D_1o_/P_1_ and a separate mutation on the terminal branch leading to P_2_), back mutation (*i.e.*, mutation on the internal branch leading to D_1_/D_1o_/P_1_ followed by a separate mutation on the terminal branch of P1 to revert back to the ancestral state), or incomplete lineage sorting (*i.e*., when a mutation occurs shortly before the divergence of D_1o_ from the D_1_/P_1_ clade, but sorts into a paraphyletic pattern). For each Diagnostic + SNP pattern, we compare the genome-wide frequency to the SNP patterns that exclude gene conversion (Diagnostic - SNPs; light blue, light red), assuming that these should occur at equal frequencies in the absence of gene conversion. That is, any hGC events will increase the number of SNP patterns consistent with hGC, while SNP patterns consistent with ILS, back mutation, or recurrent mutation should remain unaffected. Therefore, we expect the ratio of these SNP categories to be equal if no gene conversion is present, whereas a higher number of gene conversion than Diagnostic - SNP patterns would indicate hGC or homoeologous exchange.

In older allopolyploids, the number of mutations that may have occurred on the terminal branches of each diploid (D_1_ and D_2_) following their divergence from the polyploid subgenome progenitors (P_1_ and P_2_, respectively) may impact hGC diagnosis. Consider, for example, the situation where D_1_ has an autapomorphy at an otherwise invariant site; this pattern mimics the diagnostic SNP pattern expected from hGC, thus potentially leading to an overestimation of homoeologous gene conversion. Because of the possibility of these autapomorphic substitutions, the original method to detect hGC (Salmon et al., 2010) required more than one consecutive diagnostic SNP flanked by homoeoSNPs to be considered as evidence for hGC (although no flexible threshold to account for different ages of polyploids was specified). To reduce the impact of autapomorphies on inferences of hGC, we expanded phylogenetic sampling to include species closely related to the model diploid progenitors. Specifically, we include an outgroup (D_1o_ and D_2o_) for each of the diploid/subgenome clades to phylogenetically diagnose autapomorphic SNPs in each diploid (D_1_ and D_2)_, thereby allowing the identification and removal of these putative hGC sites as potential sources of error. Additionally, we included a phylogenetic outgroup to the entire genus to determine the internal branch on which these hGC-informative SNPs arose, allowing us to consider both bias in the direction of gene conversion (toward P_1_ or P_2_) and the extent to which derived alleles revert (or convert) back to their ancestral state.

This expanded phylogenetic sampling produced a set of putatively diagnostic SNPs that only includes sites where the diploid progenitor (*e.g*., D_1_) and its outgroup (*e.g*., D_1o_) both harbor derived alleles or are the only two with ancestral alleles (Figure 1B, dark red boxes). Quantification of these SNP patterns, however, does not directly measure the rate of hGC as other evolutionary phenomena could also give rise to these SNP patterns. For example, the SNP patterns in which only D_1_ and D_1o_ contain derived alleles (Figure 1B, first red box from left) could also be caused by three additional mechanisms: (1) separate, recurrent mutations on the terminal branches of D_1_ and D_1o_; (2) a mutation in the common ancestor of D_1_, D_1o_, and P_1_ that undergoes incomplete lineage sorting (ILS) to transmit the derived allele to only D_1_ and D_1o_ ; and (3) a mutation in the common ancestor of D_1_, D_1o_, and P_1_, followed by a back mutation in the P_1_ terminal branch subsequent to allopolyploid formation.

While these additional evolutionary scenarios may initially appear to unnecessarily complicate inferences of hGC, we can leverage the symmetrical properties of a phylogeny to estimate the frequency with which these scenarios affect our diagnostic SNPs. Since we expect these other evolutionary phenomena (e.g., ILS) to be distributed across branches independently of ploidy level, we can assume they produce hGC-like phenomena equally across symmetrical branches, forming a baseline estimate for each. For example, in scenario 1 (recurrent mutation), we can assume that the mutation rate on the terminal branch of D_1_ is equal to that on the terminal branch of P_1_. Thus, if we find an equal number of SNP patterns that can be explained by recurrent mutation on the D_1_ and P_1_ terminal branches (when there is also a mutation on the terminal branch of D_1o_, as explained above), respectively, we can infer that there is no hGC present in our samples. Homoeologous gene conversion would therefore be indicated by an excess of SNP patterns consistent with hGC (henceforth, “Diagnostic + SNPs”) compared to SNP patterns that can only be explained by recurrent mutation (henceforth, “Diagnostic - SNPs”). Similarly, under ILS (scenario 2), there is an equal probability that the derived alleles will be present in D_1o_/D_1_ versus D_1o_/P_1_ (but not D_1_/P_1_ due to more recent shared phylogenetic history); therefore, in the absence of hGC, we would expect the number of hGC-like SNP patterns (Diagnostic + SNPs) to equal those explained by the other ILS patterns (Diagnostic - SNPs). Finally, to estimate the influence of back mutations (scenario 3), we expect that the number of back mutations in the P_1_ terminal branch is equal to the number of back mutations in the D_1_ terminal branch. Thus, if no hGC has occurred, we would expect an equal number of Diagnostic + SNP patterns indicative of gene conversion as those where only D_1o_ and P_1_ contain the derived allele (*i.e*., Diagnostic - SNPs).

The second category of SNPs that may indicate hGC are those in which both D_1_ and D_1o_ are the only species in the phylogeny that contain the ancestral allele (Figure 1B, second red box from left). These SNP patterns may arise via only two evolutionary scenarios: recurrent mutation and hGC. Recurrent mutation would occur by a mutation in the common ancestor of D_2_/D_2o_/P_2_, followed by a recurrent mutation in the terminal branch of P_1_. Because we expect the number of mutations occurring on the terminal branch of P_1_ to be the same as those occurring on the terminal branch of D_1_, the number of SNP patterns caused by recurrent mutations in which only D_1_ and D_1o_ contain the ancestral allele (*i.e*., Diagnostic + SNPs) should be equal to the number of SNP patterns in which only D_1o_ and P_1_ contain the ancestral allele (*i.e*., Diagnostic - SNPs). Thus, any excess in the number of Diagnostic + SNPs relative to Diagnostic - SNPs would be evidence for hGC.

The above logic and scenarios are applicable and symmetrical with respect to the direction of hGC (in the present case where subgenome P_1_ overwrites P_2_ [Figure 1B, dark blue boxes]), and can be extended to situations in which the subgenomes have reciprocally ‘overwritten’ each other (Figure 1B, purple boxes). Therefore, our test for hGC not only considers the presence of SNP patterns consistent with hGC (Figure 1B, dark red and dark blue boxes), but also evaluates the abundance of these Diagnostic + SNPs relative to SNP patterns that can be explained by other evolutionary mechanisms (Figure 1B, light red and light blue boxes), providing a baseline measure for these confounding phenomena. As such, the presence of hGC can be inferred by an excess of Diagnostic + SNPs, with the absence of hGC being evidenced by equal numbers of Diagnostic + and - SNPs in our dataset.

Furthermore, this method may also be useful in identifying regions of the genome that have experienced other mechanisms of reciprocal homoeologous recombination (Figure 1B, dark purple boxes), including homoeologous exchanges, which we expect to affect larger regions of the genome and to be biased towards the telomeric regions. These reciprocal homoeologous recombination patterns are notoriously difficult to identify, however, because they result in no change in allelic dosage and because they may be artificially created by genome assembly errors, in part due to incorrect subgenome assignment.

While the aforementioned logic can be applied to genome-wide SNP counts, it is also important to consider approaches to identify particular genomic *regions* that have experienced hGC or HE, as well as the direction of these exchanges. We can use unaffected homoeoSNPs as potential ‘outer bounds’ for regions in the genome containing Diagnostic + SNP patterns where a potential hGC could have occurred. For example, by comparing the number of regions in which there are three sequential Diagnostic + SNPs versus three sequential Diagnostic - SNPs in the same direction, we can calculate the proportion of those regions that have experienced hGC using a simple D statistic.

A final consideration in the methodological logic concerns the possibility of interspecific gene flow, which can create SNP patterns similar to that of hGC, thereby complicating its detection. There are several scenarios of hybridization that may lead to similar SNP patterns as those indicative of hGC. For example, a mutation that occurs in the D_2o_ terminal branch, followed by gene flow from D_2o_ into the ancestor of D_1_, D_1o_, and P_1_ would create SNP patterns in which mutations are shared by D_1o_, D_2_, D_2o_, and P_2_ (which is a Diagnostic + SNP pattern). Likewise, mutations that occur on the common ancestor of D_2_, D_2o_, and P_2_, followed by gene flow from any of these lineages into D_1_, would create a Diagnostic - SNP pattern, thereby leading to an underestimation of the rate of hGC. Therefore, care should be taken when using these methods in systems with likely or recurrent gene flow, removing those genomic regions influenced by hybridization before interpreting results of hGC.

### No History of Interspecific Hybridization in Gossypium Diploids

Because detection of hGC using a phylogenetic SNP-based approach may be biased in the presence of introgression between diploid groups, we sought to identify potentially introgressed regions using F_DM_ statistics (39) for all possible trios of diploids in *Gossypium*. We analyzed the genome in 50 SNP windows with a sliding scale of 10 SNPs, using *Gossypioides kirkii* as the outgroup in all cases. Unsurprisingly, we found no evidence of any introgression in any of the trios tested (Supplementary Figure 1), presumably because our two diploid clades are from Central America and Africa and have remained separated by the Atlantic Ocean since their divergence 5-10 million years ago, aside from the apparently ephemeral contact 1-2 million years ago that resulted in the polyploid clade. This analysis, however, is necessary as there remains the possibility, however small, that following the migration of an A-genome propagule to the American continents, there could have been introgression from the A-genome group into the progenitor species of the polyploid D-subgenome. Evidence for introgression of rDNA and other repeated sequences has been previously suggested in *G. gossypioides* (*40–42*), although genome-wide analyses have not replicated this finding for the rest of the nuclear genome or for additional species (43) suggesting that this introgression may have been limited to *G. gossypioides*.

### Identifying Potential Regions of Homoeologous Gene Conversion in Gossypium

Because all seven polyploids in *Gossypium* diverged from the same polyploidization event (29, 33), we used the same set of diploids to identify our Diagnostic + and - SNP patterns in each polyploid species. Namely, the A-diploid progenitor (D_1_, Figure 1B) is represented by *G. arboreum* (species label A2; see Methods for genome designations); the A-diploid outgroup (D_1o_, Figure 1B) is represented by *G. longicalyx* (species labeled F1); the D-diploid progenitor (D_2_, Figure 1B) is represented by *G. raimondii* (species label D5); and the D-diploid outgroup (D_2o_, Figure 1B) is represented by *G. turneri* (species label D_1o_). All SNPs were polarized into ancestral or derived states using *Gossypioides kirkii*, an outgroup that diverged from *Gossypium* circa 6-12 million years ago (44). We initially analyzed each polyploid independently, treating the A-subgenome (*i.e*., “At” for “A-tetraploid”) and the D-subgenome (*i.e*., “Dt”) as P_1_ and P_2_, respectively (Figure 1B). Phylogenetic relationships of these species are depicted in Figure 2A.

**Figure 2:**
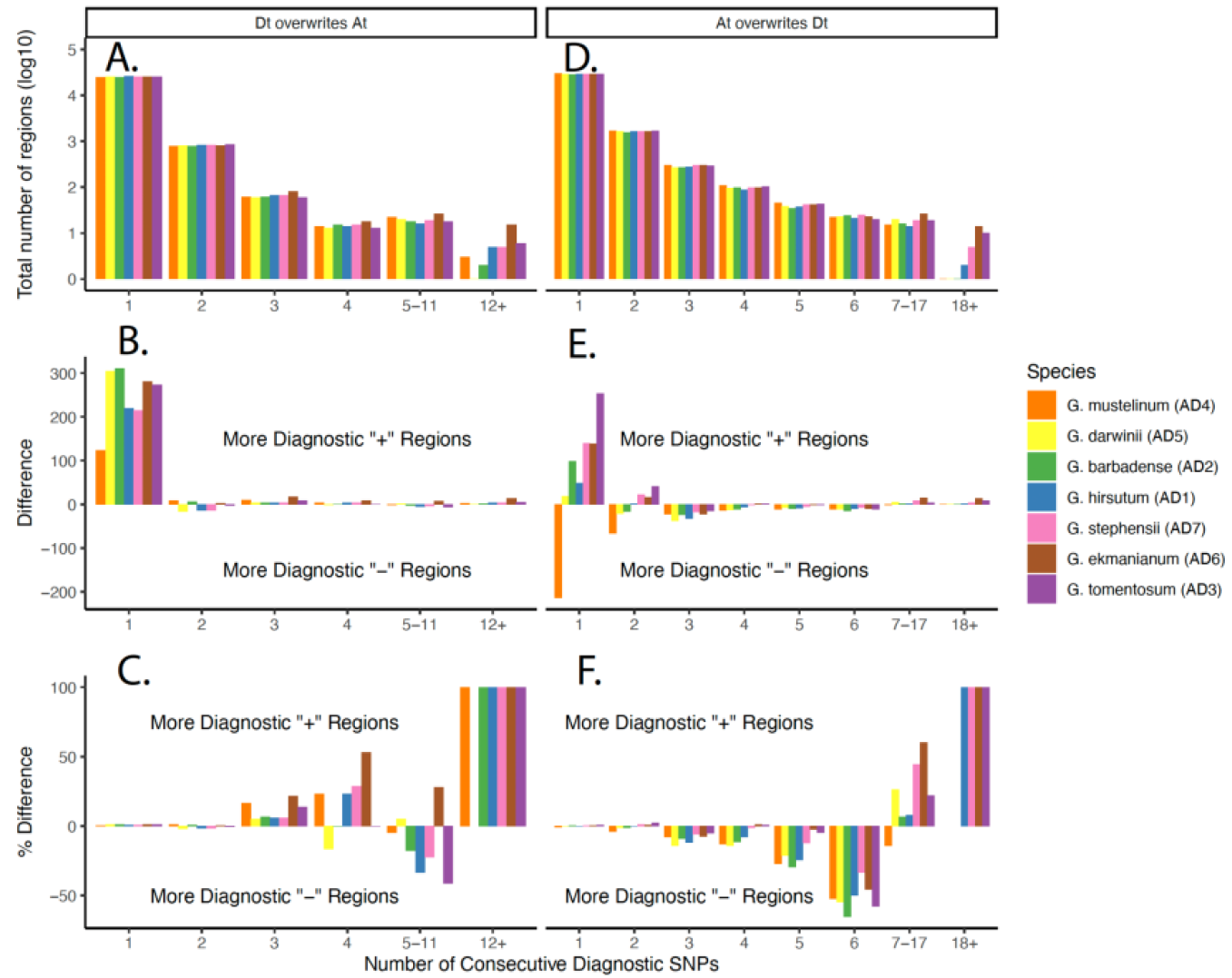
Patterns of SNPs Indicative of Homoeologous Gene Conversion and Homoeologous Exchange. Regions of the genome indicative of the Dt subgenome overwriting the At subgenome (**A**, **B**, and **C**) or the reciprocal direction (**D**, **E**, and **F**) were identified by first identifying homoeoSNPs (i.e. SNPs in which a subgenome, its most closely related diploid progenitor, and diploid outgroup all have one allele, while the other subgenome and its mostly closely related diploid progenitor and diploid outgroup have a different allele at that site). The number of Diagnostic + or Diagnostic - SNPs located between two flanking homoeoSNPs is tabulated along the x axis of each plot. (**A, D**) Total number of regions (y axis) with a given number of consecutive Diagnostic + and - SNPs (x axis). (**B, E**) The difference in the number of regions with Diagnostic + compared to Diagnostic - SNPs. Positive values indicate more regions with Diagnostic + SNPs, negative values indicate more regions with Diagnostic - SNPs. (**C, F**) The percent difference between the number of regions with Diagnostic + or - SNPs, calculated as (# Diagnostic + regions / total number of regions). Each polyploid is represented by a different color in the bar graph, shown at right.

For each polyploid, we were able to identify over 1 million homoeoSNPs that distinguish the two homoeologous subgenomes (ranging from 1.01M in AD2 to 1.05M in AD1; Supplementary Figure 2). While the number of homoeoSNPs is considerably lower than previously reported (∼25M, (45)), we note our strict requirements for homoeoSNP definition. That is, homoeoSNPs were only inferred when all members of the D5/D10/Dt clade (where “Dt’’ refers to the D-subgenome in the tetraploid) shared a SNP that was different from the SNP shared by all members of the F1/A2/At clade (Figure 1B, Grey Box). As expected, more derived alleles were found at the base of the D clade compared to the A clade (Supplementary Figure 2) due to the relatively more recent divergence of D5/D10/Dt (∼1.76 Mya (43)) versus F1/A2/At (∼4 Mya (37)). Additionally, there were more derived mutations present on the terminal branches of both subgenomes of all polyploids (with the exception of the At subgenome of *G. mustelinum* (AD4)) compared to their respective diploid progenitor (Supplementary Figure 2), consistent with previous findings (34) and which may be a result of the masking effects of allopolyploidy that reduces the fitness consequences of deleterious alleles (46) or from unequal evolutionary rates between the subgenomes as compared to their diploid relatives (47).

### Homoeologous Gene Conversion Where Dt Overwrites At

For each polyploid, we identified between 24,900 and 26,000 regions that were flanked by homoeoSNPs and also contained at least one Diagnostic +/- SNPs (Figure 2A) that would be consistent with the Dt subgenome overwriting the At subgenome. These regions ranged in size from a single nucleotide (*i.e*., two homoeoSNP had a single base pair between them, and that base pair showed a SNP pattern consistent with a +/- Diagnostic SNP) to over 576kb, with a strong bias toward smaller regions. Only a single Diagnostic +/- SNP was identified in most regions (Figure 2A), and the number of identified regions decreased as the number of consecutive Diagnostic +/- SNPs increased. The total distribution of Diagnostic + SNPs compared to the distribution of Diagnostic - SNPs showed no statistical significance (*p* value > 0.99 for all species, two-sided Kolmogorov–Smirnov test); thus, for graphical clarity, we combined all regions in which the number of Diagnostic +/- SNPs in all seven species ranged from 5-11 (5 being the smallest bin size in which some species had fewer than 10 regions, and 11 being the largest number of consecutive Diagnostic - SNPs, although these are arbitrary cutoff points used only for graphical clarity). The region with the largest number of Diagnostic - SNPs (*i.e*., those consistent with ILS or recurrent mutation, but not hGC or HE) in all species contained 11 SNPs, while the region with the highest number of Diagnostic + SNPs contained 150 SNPs. Interestingly, these higher numbers of Diagnostic + regions were clustered at the terminus of chromosome D5_01 in six of the seven polyploids in our analysis, indicating a homoeologous exchange event rather than a hGC event (discussed below).

Because the Diagnostic + SNP patterns can reflect evolutionary processes other than homoeologous gene conversion, we compared the proportion of Diagnostic + to Diagnostic - SNPs in each polyploid species to assess the putative rate of hGC. If hGC has historically occurred in any of these lineages, we expect to see a marked excess in the proportion of Diagnostic + SNPs relative to Diagnostic - SNPs. In contrast to our *a priori* expectations based on earlier hGC assessments, we see no enrichment of Diagnostic + SNP patterns (relative to Diagnostic - SNPs) for those regions containing four or fewer consecutive diagnostic SNPs, either when comparing the total number of SNPs within regions (Figure 2B) or in the proportion of SNPs that exhibit Diagnostic + SNP patterns compared to the total number of SNPs (Figure 2C). Interestingly, we see a higher fraction of regions that contain Diagnostic - compared to Diagnostic + SNPs in several of these categories, potentially indicating a higher rate of recurrent mutations on the terminal branches of the polyploids as compared to the terminal branches of the diploid progenitors. For regions that only have a single Diagnostic SNP (Figure 2B), we see a higher number of Diagnostic + SNPs for all species indicating that a small amount of hGC may be occurring that predominantly affects small regions of the genome; however, because we only see 100-300 excess regions out of a possible 10,000 total regions, this difference is not statistically significant and should not be interpreted as evidence for hGC taking place.

### Homoeologous Gene Conversion Where At Overwrites Dt

For those SNP patterns that are consistent with hGC in the opposite direction, (*i.e*., At overwriting Dt; Figure 2D), we see a slightly higher number of regions (29,000–32,000), as described above for potential hGC, presumably due to a more distantly related diploid outgroup on this side of the phylogeny (*G. longicalyx*) compared to the other (*G. turneri*), allowing for a larger proportion of sites with SNP patterns caused by recurrent mutation to pass filtering. The broad size patterns of these SNPs across the genome are similar to those described above (ranging in size form 1bp to 622kb), where regions in which one Diagnostic + or - SNP is contained within a single region flanked by homoeoSNPs is the most common. The region with the highest number of Diagnostic - SNPs contained 11 SNPs, and the total distribution of Diagnostic + compared to the distribution of Diagnostic - SNPs showed no statistical significance (*p* value > 0.88 for all species, two-sided Kolmogorov–Smirnov test). Therefore, we combined some groups together for graphical clarity depending on the total number of sites present in each species; namely, all species had at least 10 sites with 6 or fewer consecutive Diagnostic + SNPs, and the highest number of consecutive Diagnostic - SNPs in any species was 17, so we combined groups of size 7-17. The difference in the highest number of consecutive Diagnostic + to Diagnostic - SNPs is different for the two directions of potential hGC or HE (17 when Dt overwrites At, 11 when At overwrites Dt) due to the difference in the phylogenetic relatedness of the diploid outgroups in the two clades and is an important aspect of choosing samples for repeating this analysis in other systems (see Discussion). The regions with the largest number of Diagnostic + SNPs in any species contained 105 (chromosome D5_05 in *G. tomentosum* (AD3)) and 76 (chromosome D5_12 in *G. ekmanianum* (AD6)) such SNPs; however, because these regions are restricted to a single species and do not occur at the termini of the chromosome arms, as would be expected under HE events, it is difficult to differentiate whether these patterns are caused by real hGC events or are due to artifacts, such as alignment errors or genome assembly errors. Notably, *G. ekmanianum* and *G. tomentosum* contained considerably higher numbers of regions with elevated numbers of Diagnostic + SNPs (Figure 2F; Figure 3), indicating that these genome assemblies (or alignments) are of poorer quality than those of the other polyploids in the clade or that there may be a temporal and lineage-specific variation in the rate and occurrence of HE and/or hGC in *Gossypium*. When comparing the difference in the number of regions with Diagnostic + or Diagnostic - SNPs indicative of At overwriting Dt (Figure 2E), we again see similar patterns as described above (Figure 2B). Congruent with our prior analysis, we see no evidence of enrichment in Diagnostic + SNPs (versus Diagnostic - SNPs) for regions of size bins 1 or 2, although in most species, a small number of regions have a single Diagnostic + SNP more than regions that have a single Diagnostic - SNP, with the exception of *G. mustelinum*. As mentioned above, because this small difference is tabulated from a total of over 10,000 regions, this is not statistically significant and should not be interpreted as evidence of hGC taking place.

**Figure 3:**
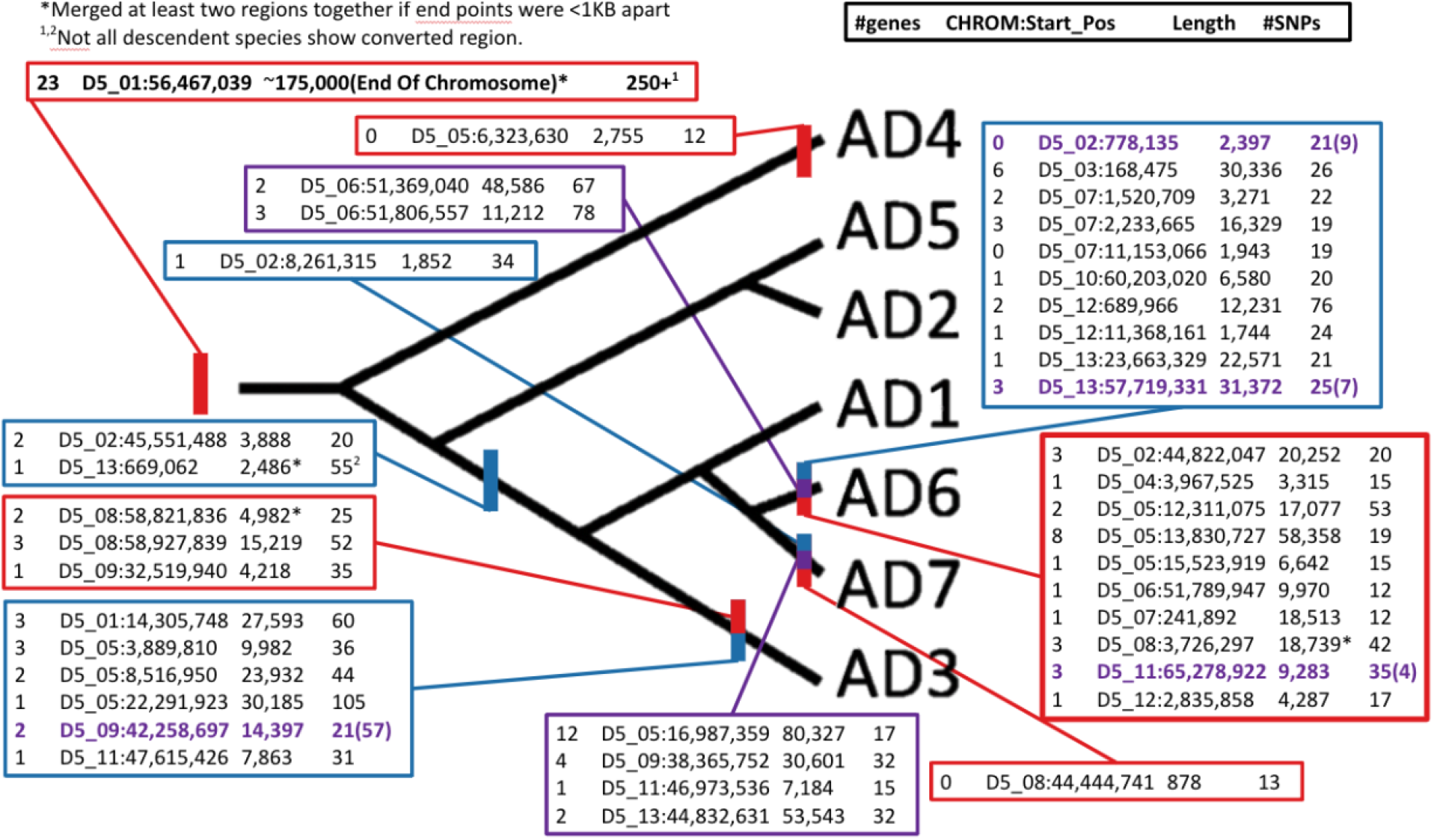
Evolutionary Timing of Potential Homoeologous Gene Conversion or Homoeologous Exchanges in Gossypium Phylogeny. Using only the regions which were longer than the longest region consistent with patterns of Diagnostic - SNPs (*i.e*., at least 12 SNPs when Dt overwrites At [outlined in red], and at least 18 SNPs when At overwrites Dt [outlined in blue], and at least 4 SNPs where reciprocal hGC has occurred [outlined in purple]), we can map each of these events to the *Gossypium* phylogeny using parsimony. For each line, the four columns represent the number of genes in the region, the chromosome/starting point of the region, the length of the region, and the number of consecutive Diagnostic + SNPs in that region, respectively. If any region included more than three SNPs indicative of reciprocal hGC, the text of that region is purple and the number of reciprocal SNPs is included in parentheses. The phylogeny does not represent scaled evolutionary distances or divergence between species, only the relative relationships as inferred from previous analyses (34, 35).

### Potential Gene Conversion Events Where Diagnostic + SNP Tracts are Longer than Diagnostic - SNP Tracts

In six of the seven allopolyploids (*i.e.*, all except *G. darwinii*, AD5), there was at least one genomic region that contained more consecutive Diagnostic + SNPs than the longest track of consecutive Diagnostic - SNPs observed in any species. In total, we found 19 regions of this type that are consistent with the At subgenome overwriting the Dt subgenome (Figure 3). Two of these regions were found in more than one species (with identical locations for flanking homoeoSNPs) and occur in parsimonious positions along the polyploid phylogeny. One region found on chromosome D5_02 in *G. hirsutum* (AD1), *G. tomentosum*, (AD3), *G. ekmanianum* (AD6), and *G. stephensii* (AD7) is 3,888 base pairs long, contains twenty consecutive Diagnostic + SNPs, and partially overlaps with two genes. Because these four polyploids form a monophyletic clade, it is more likely that this region reflects an event in the common ancestor of these four species following their divergence from the other three polyploid species (*G. mustelinum* (AD4), *G. barbadense* (AD2), and *G. darwinii* (AD5)) rather than an error in genome alignment. A second region on chromosome D5_13 was 2,458 base pairs in length and partially overlaps one gene. This region, however, was found in *G. tomentosum* (AD3), *G. ekmanianum* (AD6), and *G. stephensii* (AD7), which form a monophyletic clade only with the inclusion of *G. hirsutum* (AD1). Interestingly, however, we find evidence that this region in the *G. hirsutum* genome may have experienced introgression from *G. barbadense* (AD2) (Supplementary Figure 3), either as a result of historical natural introgression between these two species or as a result of the known intentional introgression during crop domestication and improvement (48). Each of the remaining 17 regions that contained long stretches of Diagnostic + SNPs (affecting 0-6 genes each, for 32 total) were confined to a single species, potentially revealing recent hGC and/or HE events, although we cannot rule out genome assembly errors and/or species-specific errors in genome alignments. The latter suggestions (*i.e*., error) may be more reasonable in this case, as the species-specific observations occurred in only three species (*i.e*., *G. tomentosum* (AD3), *G. ekmanianum* (AD6), and *G. stephensii* (AD7)), most of which were detected in only *G. ekmanianum* (10 regions, 19 genes) and *G. tomentosum* (6 regions, 12 genes). Given the general paucity of evidence for hGC and HE in the rest of our analyses, we posit that these regions likely represent artifacts rather than true hGC or HE events.

Similarly, we also found 16 regions consistent with the Dt subgenome overwriting the At subgenome in which the number of Diagnostic + SNPs was higher than any region with consecutive Diagnostic - SNPs, and only one of these regions was shared by more than one species. This single region was shared by all species except *G. darwinii* (AD5) and affected the terminal 175 Kb (23 genes) of chromosome D5_01. The number of putatively converted SNPs in this region varied from 120-261 among species (see below), but the ‘starting point’ (*i.e*., the homoeoSNP most distant from the chromosome terminus) is consistent in all six polyploids, resulting in the complete conversion of 23 genes. We explore this region in detail below. The remaining 15 regions (affecting 0-8 genes each) were all present in a single species, again with with the majority found in *G. ekmanianum* (10 regions, 24 genes) and the remainder found in *G. tomentosum* (3 regions, 6 genes), *G. stephensii* (1 region, 0 genes), and *G. mustelinum* (1 region, 0 genes).

### A Single, Shared Homoeologous Exchange Event Occurred Shortly After Polyploidization

The region described above, involving the final 175 Kbp and 120-261 SNPs on the end of chromosome D5_01, is indicative of a HE event that results from recombination between the subgenomes of an allopolyploid in which the Dt subgenome has overwritten the At subgenome. Typically, HE events affect the entire terminus of a chromosome past the recombination breakpoint, and this region is then fragmented by homologous recombination in subsequent generations in an analogous way that blocks of introgression are broken up by recombination. This region, however, contains few SNPs that are indicative of recombination breaking up the region in any of the polyploids, suggesting that the initial recombination event leading to the HE either occurred in a very small population, perhaps immediately following the bottleneck that is typically associated with allopolyploid formation, or has experienced strong positive selection, leading to the fixation of the HE in the ancestral population before the first speciation event occurred. Supporting this, we note that in the six polyploids that show evidence of this HE event, all displayed the same initial recombination point and all contained few places of homologous recombination that broke up this HE event into smaller regions. Notably, one species in our analysis (*G. darwinii*, AD5) failed to show evidence of this HE event that is shared by all other polyploid species; however, further inspection suggests this may be an assembly artifact (Figure 4). Inspection of genome alignments between all of the polyploids and their diploid progenitors revealed a gap in the assembly at the ends of both chromosomes D01 and A01 in *G. darwinii*, suggesting that the initial lack of evidence for this HEe may be the result of inter-subgenomic sequence similarity (due to the HE) impeding the accurate assembly of homoeologous chromosomes A01 and D01 in *G. darwinii*. Interestingly, this region also exhibits missing sequences in some of the other allopolyploid genomes (*e.g., G. barbadense* (AD2)). These gaps, however, did not span the entire HE region, and thus did not inhibit our ability to detect the HE event (Figure 4B).

**Figure 4:**
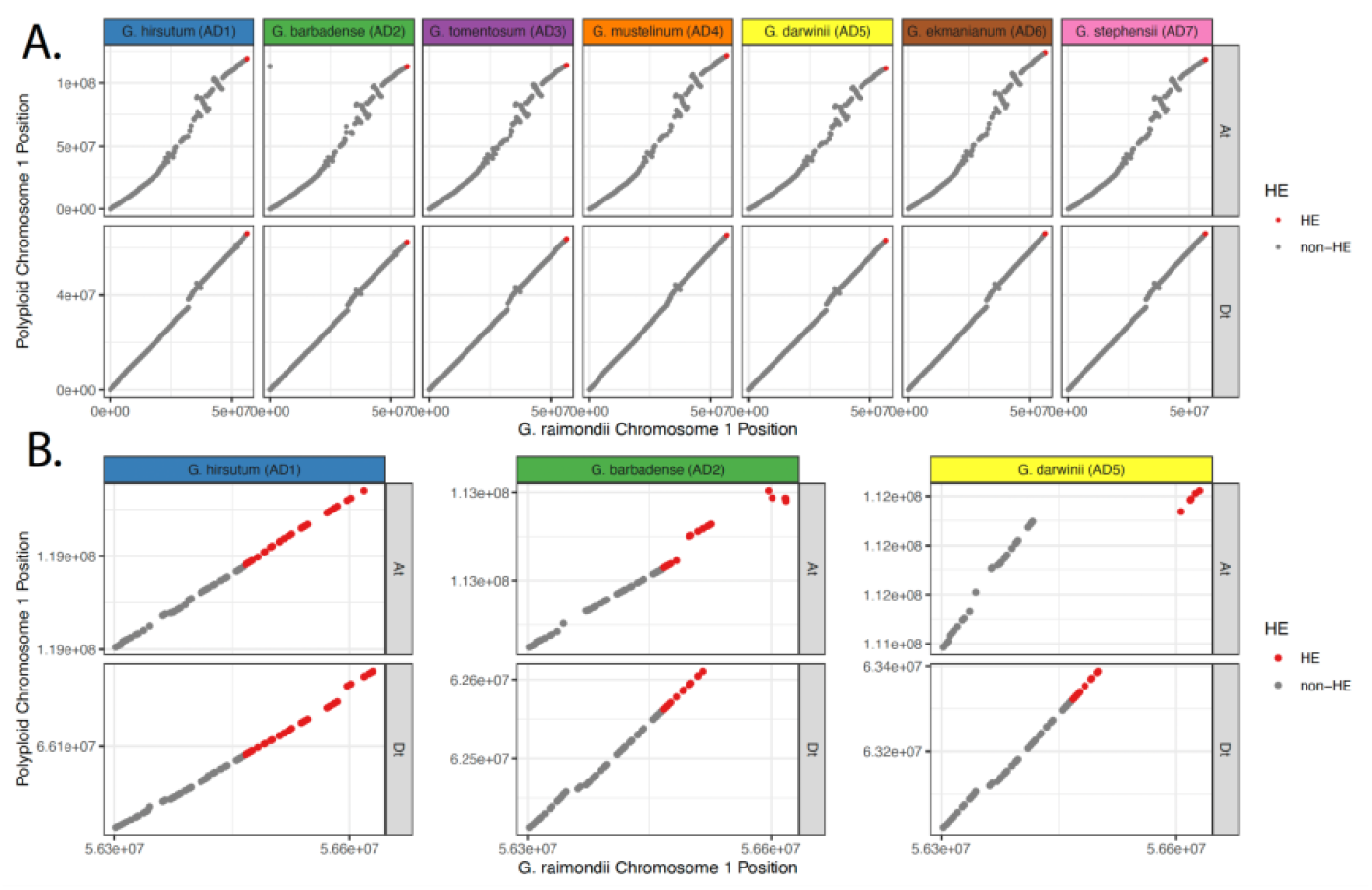
Homoeologous Exchange Event on Chromosome 1 Locates To A Gapped Region in All Polyploid Genome Assemblies. **(A)** Dotplot showing the whole-genome alignment between each subgenome (top: At subgenome; bottom: Dt subgenome) aligned to the *G. raimondii* genome. Anchors identified by Anchorwave are shown here. Colors of the facet labels are consistent with the color scheme used through the other figures. **(B)** Dotplot of the region that has experienced a homoeologous exchange event. Regions that are inferred to have experienced the HE event are shown in red, and regions that did not experience the HE event are shown in grey. In *G. hirsutum* (left, Blue), accurate assembly of both subgenomes allowed for the accurate identification of the HE event. In *G. barbadense* (middle; Green), a gap in the At subgenome (top) and incomplete assembly of the telomeric region of the Dt subgenome (bottom) did not hinder HE identification, even though the HE is the likely cause of the assembly difficulties, as the regions directly flanking the initial recombination event (i.e. where the grey dots meet the red dots) is still intact in both subgenomes. In *G. darwinii* (right, Yellow), a large gap in the At subgenome (top) and incomplete assembly of the telomeric region of the Dt subgenome (bottom) led to the inability to positively identify this region as an HE since there is no vertical region in which there are two anchors shown in red.

### Reciprocal hGC and HE

Finally, we evaluated SNP patterns consistent with reciprocal gene conversion where we simultaneously observe consecutive At SNPs on the Dt chromosome and vice versa for the same region (Figure 1B, dark purple boxes). Although the mechanism of hGC is inherently unidirectional, it is possible that hGC may occur in different individuals in different directions at the same time, thus leading to hGC in both directions. Additionally, these SNP patterns may be caused by HEs (which are inherently reciprocal) or by errors in genome assembly where the subgenome assignment is incorrect. While it is difficult to estimate the expected frequencies of counterbalancing SNP patterns that could be explained equally well by recurrent mutations, ILS, and/or back mutations (as we created for the previous gene conversion analyses), we found these SNP patterns to be rare, indicating that reciprocal hGC or HE is not likely to have occurred in any *Gossypium* allopolyploid. In four of the polyploids (*G. hirsutum* (AD1), *G. barbadense* (AD2), *G. mustelinum* (AD4), and *G. darwinii* (AD5)), we found three or fewer regions with two consecutive SNPs indicative of reciprocal gene conversion (and never saw more than two consecutive SNPs fitting this pattern). We observed longer regions with putative reciprocal gene conversion SNP patterns in the other three polyploids (*i.e*., *G. tomentosum* (AD3), *G. ekmanianum* (AD6), and *G. stephensii* (AD7)), although we approach these with caution. In *G. tomentosum*, we identified a single region that contained 57 consecutive SNP patterns consistent with reciprocal gene conversion, and a second region containing only two neighboring SNPs congruent with reciprocal hGC. We note, however, that *G. tomentosum* was among those species mentioned above where possible assembly errors artifactually generated observations of hGC, and that these assembly errors could produce SNP patterns consistent with reciprocal hGC. Likewise, both *G. ekmanianum* (19 regions, 3-78 SNPs) and *G. stephensii* (4 regions, 15-32 SNPs) exhibited SNP patterns consistent with reciprocal gene conversion, and while these regions were among the largest discovered in our analysis (between 7 KB - 80 KB, affecting 1-12 genes each), we note that these genomes were also among those suspected of assembly artifacts, similar to *G. tomentosum*.

### A Direct Comparison of Quartet-Based Methods

While the primary goal of this study is to develop an updated method to detect hGC or HE events while explicitly accounting for other evolutionary processes that may create similar SNP patterns (*e.g*., ILS, recurrent mutation), we also tested the classical ‘quartet’ approach on a genome-wide basis. To do this, we tabulated the number of Diagnostic + SNPs (Figure 1A; *i.e*., SNP patterns that may be created via hGC or autapomorphic mutation on a diploid terminal branch) as well as SNPs that are analogous to our Diagnostic - SNPs in the seven-taxon test (*i.e*., SNP patterns that may be created via autapomorphic mutation on a terminal polyploid subgenome branch). Given no instances of hGC or HE, we expect that these distributions should be identical, and any deviations in which Diagnostic + SNP patterns are observed more frequently than Diagnostic - SNP patterns would suggest that hGC or HE may play a role in shaping the overall SNP patterns in the genome.

Using the quartet approach, we found nearly three orders of magnitude more regions with SNP patterns that fall into our Diagnostic + and - categories (2.14M [AD2] - 2.22M [AD1]) relative to our 7- taxon test (52.2K [AD2] - 54.7K [AD1]), likely due to the less stringent SNP thresholds in the quartet test. Thus, a higher number of these SNPs are likely due to autapomorphic mutations, recurrent mutations, or back mutations. The distribution of consecutive Diagnostic + SNPs was much broader than our 7-taxon test, with the highest number of consecutive Diagnostic + SNPs being 347 (the HE region on chromosome 1 in AD4) and the highest number of consecutive Diagnostic - SNPs between flanking homoeoSNPs being 2,140 (chromosome 12, AD5). For hGC or HE occurring in either direction, we found that regions with only a single Diagnostic + or - SNP occur most frequently (Figure 5A,B), and that number of regions decreases as the number of consecutive SNP patterns increases. Additionally, we find that the distribution becomes increasingly dissimilar between species as the total number of consecutive SNPs increases, indicating that the upper tail of these distributions may be noisy and produce unreliable or inconsistent results.

**Figure 5:**
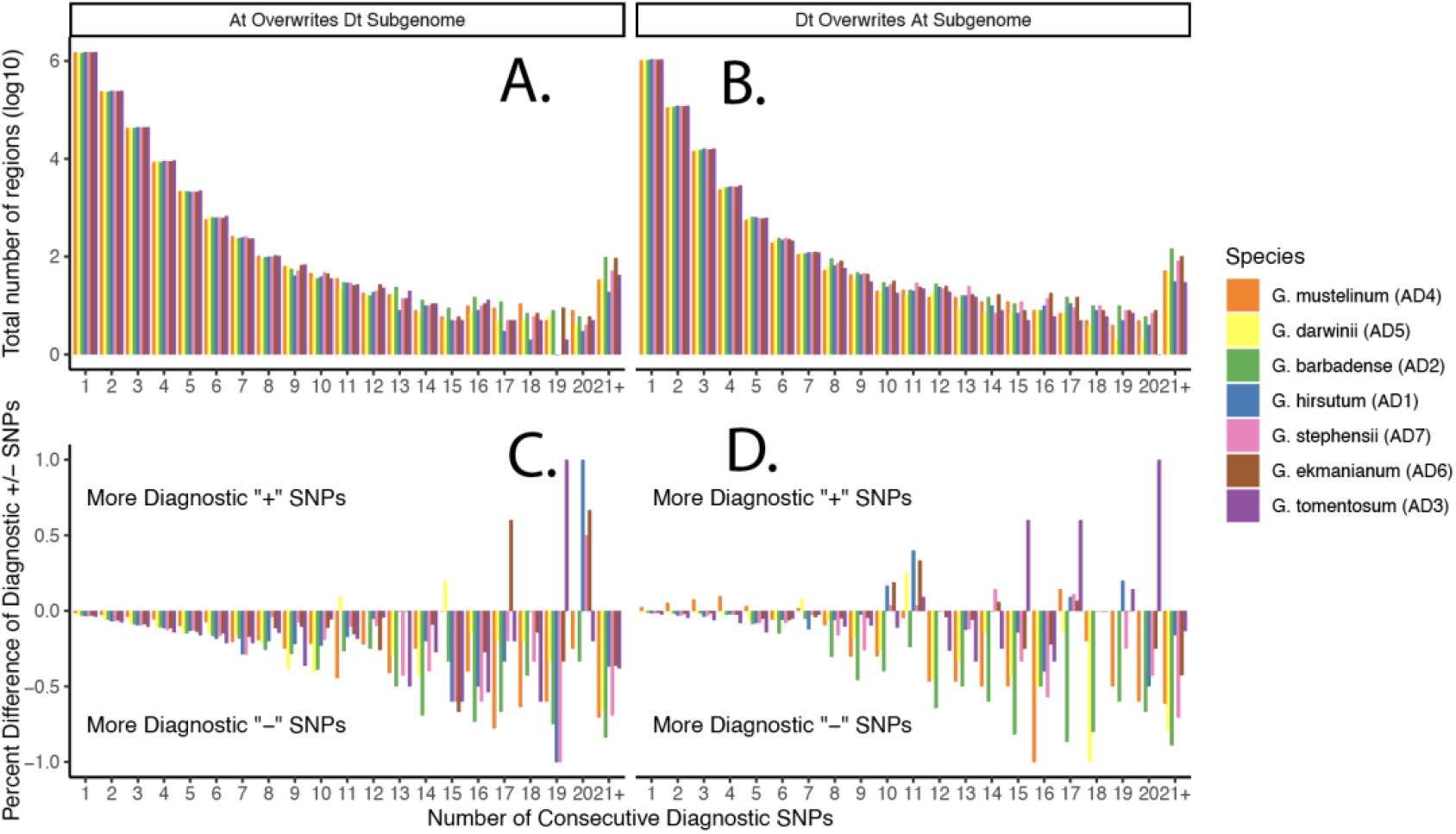
Classical “Quartet” Methods Fails to Detect Homoeologous Gene Conversion and Homoeologous Exchanges. Regions of the genome indicative of the Dt subgenome overwriting the At subgenome (**A**, **C**) or the reciprocal direction (**B**, **D**) identified using a modification of the classical ‘quartet’ method. Regions were identified by first identifying homoeoSNPs (i.e. SNPs in which a subgenome and its most closely related diploid progenitor have one allele, while the other subgenome and its mostly closely related diploid progenitor have a different allele at that site). The number of Diagnostic + or Diagnostic - SNPs located between two flanking homoeoSNPs is tabulated along the x axis of each plot. (**A, B**) Total number of regions (y axis) with a given number of consecutive Diagnostic + and - SNPs (x axis). (**C, D**) The percent difference between the number of regions with Diagnostic + or - SNPs, calculated as (# Diagnostic + regions / total number of regions). Positive values indicate more regions with Diagnostic + SNPs, negative values indicate more regions with Diagnostic - SNPs. Each polyploid is represented by a different color in the bar graph, shown at right. Although the classical ‘quartet’ method only required two consecutive Diagnostic + SNPs to be inferred as hGC, we compare the total number of regions with the same number of Diagnostic + or - SNPs to infer hGC events.

When comparing the distribution of Diagnostic + to Diagnostic - SNPs indicative of the At subgenome overwriting the Dt subgenome through hGC or HE (Figure 5C), we find that the total distribution of Diagnostic + SNPs is statistically dissimilar from that of Diagnostic - SNPs for all species (p < 2.14e-17 for all species, two-sided Kolmogorov-Smirnov test); however, because we find that an excess of Diagnostic - SNPs in every species and for every category of SNP, this difference is not due to the presence of hGC or HE. For potential hGC or HE events in the opposite direction (*i.e*., in which the Dt subgenome overwrites the At subgenome), we find similar, though not as striking, patterns with the exclusion of AD4. These distributions have varying statistical significance in each species (*p* = 1.8e-9 in *G. mustelinum*; *p* = 1.8e-7 in *G. tomentosum*; *p* = 3.4e-3 in *G. stephensii*; *p* = 3.9e-3 in *G. hirsutum*; *p* = 0.256 in *G. barbadense*; *p* = 0.413 in *G. ekmanianum*; *p* = 0.664 in *G. darwinii*; two-sided Kolmogorov-Smirnov). In *G mustelinum* (AD4), we find a slight excess of Diagnostic + SNP regions of size 1-5, but all other species had a deficit of Diagnostic + SNP regions of this size. For some of the larger values of consecutive SNPs, we see that some species have an excess of Diagnostic + SNP regions, but there is no clear trend for hGC or HE in either direction, including the previously described HE event on Chromosome 1. Thus, we conclude that quartet-based methods are unreliable in identifying either small regions of homoeologous gene conversion or larger homoeologous exchanges in allopolyploid cottons.

## DISCUSSION

Genomes of polyploid species are notoriously difficult to analyze, due in no small part to the physical interactions between duplicated chromosomes in meiosis that lead to homoeologous exchanges (HEs) or homoeologous gene conversion (hGC). Here, we developed a more robust and analytical framework for identifying the regions of an allopolyploid genome that have experienced these interactions, using a monophyletic group of seven allopolyploid cotton species with some previously described instances of hGC, but no documented cases of HEs, combined with five genomes of closely related diploid species. While previous analyses using ‘quartet’-based approaches (*i.e*., those using only the two subgenomes of an allotetraploid and a closely related diploid to each subgenome) estimated that between 1-7% of genes in allopolyploid *G. hirsutum* and *G. barbadense* have experienced hGC (8, 23), our analytical method reveals that this is a vast overestimation, suggesting that two and zero genes have potentially been influenced by hGC in *G. hirsutum* and *G. barbadense*, respectively. In the other five allotetraploid species in the clade, nearly 100 genes may have experienced hGC in total, but because nearly all of these events are found not to be shared between any species (even between extremely closely related species, as is the case with *G. ekmanianum* and *G. stephensii*), it is difficult to rule out assembly or alignment artifacts as the cause of these patterns. In total, our results suggest that traditional ‘quartet’-based methods of inferring hGCs may be unreliable and produce dramatic overestimates of the rates of HE/hGC.

We also present the first evidence of an HE event within cotton genomes, affecting the terminal 175 kb (and 23 genes) of chromosome 1 in six of the seven genomes analyzed (although the seventh species has a poorer assembly at this region in chromosome 1 of both subgenomes, indicating that it has also likely experienced this HE event). This is a notable HE event, as the two-fold difference in size between homoeologous chromosomes has generally been thought to restrict the pairing behavior of homoeologs during meiosis. As such, no multivalent chromosome pairing behavior has been observed in any of the allotetraploid cotton species. In fact, it has long been established (49, 50) that intergenomic A genome x D genome diploid hybrids in Gossypium exhibit only about six bivalents (of the 13 possible), whereas both natural and synthetic allopolyploids have nearly complete homoeologous pairing (50). Moreover, chromosome pairing in synthetic allopolyploids is limited almost entirely to homoeologs. Thus, as noted by (31) and others, the restriction of pairing to homologous chromosomes that we see today most likely existed even at initial allopolyploid formation.

The fact that we see very low rates of HE correlated with low rates of hGC is probably not coincidental, as the same meiotic machinery and pathway that generates HEs is also responsible for generating hGC. Thus, while the end result of hGCs and HEs may be dissimilar in the direct meiotic products, their long-term genomic signatures may be difficult to distinguish from each other, particularly after several generations where regions of HEs can be broken up via homologous recombination. Thus, further development of methods to identify hGC is warranted in species with documented examples of HEs, and with attention to distinguishing hGC from HE. Although the SNP patterns in regions that have experienced HE should be identical to those that have experienced hGC, other attributes of these sites not explored here (for example, analyzing the distribution of hGC or HE tract length sizes compared to the distribution of haplotype block sizes genome-wide, or through the direct tracking of recombination events through the use of ancestral recombination graphs (51)) offer a potential opportunity to distinguish these intertwined phenomena. Analytical methods for population-level processes in allotetraploids (*e.g*., demographic history, interploidy introgression) has received recent attention (52, 53), and we suggest that studying hGC and HE at the population level may offer additional insights that are not possible with our phylogenetic SNP-based approach.

While many studies have suggested that hGC could be a pathway for allopolyploid genomes to overcome genetic incompatibilities or create new haplotypes on which selection can act (54–56), our results suggest that because these studies are based on the ‘quartet’ approach, their interpretation of identifying *bona fide* hGC regions should be treated with caution. While we do not disagree that, conceptually, hGCs can create novel haplotypes that might be visible to selection, it is important to robustly and systemically test this hypothesis before concluding that a particular region of an allopolyploid genome has experienced hGC and that selection has acted upon these regions. Ultimately, the detection of hGC and inferences of selection are attempts to attribute evolutionary processes to explain a pattern observed in data; as we show here, however, multiple evolutionary processes may generate the same patterns of data.

Although our current method is unable to identify with certainty which regions of the genome have experienced hGC (only the proportion of each SNP pattern described above that are overrepresented compared to their non-hGC counterparts), it is important to note that other meiotic recombination patterns, such as double crossovers between homoeologous chromosomes, may produce SNP patterns that appear similar to longer hGC tracts that may not affect a portion of a chromosome arm in the same way that homoeologous exchanges do. However, because the typical tract length of hGCs or double crossovers involving homoeologous chromosomes is not known, it is not possible to distinguish these two processes, which remains a topic for future work, presumably in a system that has more easily identifiable regions that have experienced hGC, HE, and double crossovers. Analytical methods to identify the frequency of occurrence and the allelic segregation patterns of double crossovers is an active area of research in autopolyploids (57, 58), but extending these methods to allopolyploids that experience HEs or hGC has received little attention.

Extending this analytical pipeline to other allopolyploid systems will be of interest to others, so it is germane to consider some of the necessary criteria for taxon selection. The most important aspect of choosing species for this analysis is to identify those with minimal amounts of gene flow, so that the SNP patterns are not influenced by this inherently homogenizing effect. The species in *Gossypium* used here are probably an outlier in this respect, as most diploid progenitors to an allopolyploid are probably not distributed in different hemispheres separated by an ocean. Additionally, the availability and relative divergence of diploid outgroups matters, as it influences the total number and relative symmetry of SNP sites. We saw a marked increase between the total number of SNP sites in the A lineage that were attributable to either ILS, recurrent mutations, or back mutations, compared to D lineage. This likely is due to the phylogenetic position of the diploid outgroups, where the diploid outgroup for the A lineage (F1) was considerably more distant to the polyploidy subgenome than the D lineage outgroup (D10). Hence, there are more opportunities for recurrent mutations and back mutations to occur on the terminal branches of F1, and either of the terminal branches of A2 or the At subgenome. Although we did not directly test this, this observation suggests that ILS was responsible for a comparatively small portion of the SNPs used here and that the majority of the Diagnostic - SNPs may be caused by recurrent or back mutations. Thus, when choosing a diploid outgroup, there is a tradeoff between choosing a species that is distantly related enough to minimize the amount of ILS, with one that is close enough to minimize the amount of recurrent or back mutation. Finally, we note the requirement for high-quality genome assemblies for allopolyploids (and the increase in interpretive possibility from including more than one) as well as genomes for both model diploid genome donors and outgroups to these as well as to the system overall. Although these are relatively stringent methodological requirements, the number of high-quality genomes continues to increase in recent years, and the number of genomic tools such as whole genome alignment algorithms designed specifically for the complexities of plant genomes is under active development (59). Thus, the application of methods such as those presented here may provide additional insight into questions addressing the genomic location, rate, and tempo of hGC and HE events, as well as their consequences for selection and adaptation in allopolyploids.

## MATERIALS AND METHODS

### Data and Genome Alignments

Nomenclature for Gossypium species and their genomes has been standardized (Wang et al., 2018) and is used here. Specifically, the following designations are used for A-genome diploids (A1 = *G. arboreum*) and D-genome diploid species (D5 = *G. raimondii*, D10 = *G. turneri*), and F-genome diploids (F1 = *G. longicalyx*). In addition, the two co-resident genomes in each allopolyploid species are indicated by symbols representing their origin (A or D) along with the subscript t, for tetraploid, to distinguish them from their diploid counterparts. Genome sequences for seven allotetraploid genomes (*Gossypium hirsutum* (AD1; accession Bar32; (60))*, G. barbadense* (AD2)*, G. tomentosum* (AD3)*, G. mustelinum* (AD4), and *G. darwinii* (AD5), (34), and *G. ekmanianum* (AD6) and *G. stephensii* (AD7) (35)), two model diploid progenitors (*G. raimondii* (38) and *G. arboreum* (61)), an outgroup to each diploid/subgenome clade (*G. turneri* (38) and *G. longicalyx* (37)), and an outgroup to the entire genus (*Gossypioides kirkii* (44)) were downloaded from CottonGen (62). Any scaffolds or contigs not anchored to the pseudochromosomes of each assembly were removed. Genomes for the polyploids were split by subgenome and independently aligned to a diploid reference genome using AnchorWave (63) (last accessed: July 6, 2022), with annotations ported to each genome using gsnap (64). We used the proalign function within AnchorWave while allowing for the possibility of relocation variation, inversion, or chromosome fusion using flags -R 1 - Q 1 and -m 0.

Alignments of paralogous genomic regions, copy number variants, and/or presence/absence variants can easily be misinterpreted as phylogenetically discordant regions and, hence, gene conversion or homoeologous exchange events. Therefore, we performed strict filtering to remove these regions from our whole genome alignments. Because the size of the genomes of the diploid species of interest vary by nearly twofold (not including the fold-difference due to polyploidy *per se*), we took extra precautions to ensure that our analyses were not influenced by alignment errors. In particular, we aligned every genome to the smallest (*G. raimondii*, 885 Mbp) and largest (*G. arboreum*, 1,700 Mbp) genomes sampled within *Gossypium*. Pairwise alignment files were converted to gVCF files using the MAFToGVCFPlugin tool from the Practical Haplotype Graph project (65) and all gVCF files were collated into a multi-sample vcf using bcftools (66). Any sites including indels or non-biallelic sites were filtered out using vcftools (67). We also ensured that, for a given diploid species or polyploid subgenome, we excluded any regions that mapped to different loci between the two diploid reference genomes using custom python scripts. Scripts for all alignments and data filtration are available on Github (https://github.com/conJUSTover/GeneConversion), and raw alignment and filtered VCF files are available on Figshare (DOI: 10.25422/azu.data.24512896).

### Detecting HomoeoSNPs and Potential Converted Regions

To detect SNP patterns that are either indicative of potential hGC or HE events (Diagnostic + SNPs) or to create the null expected number of these SNP without invoking processes of hGC or HE (Diagnostic - SNPs), we developed a custom Python script that parses a VCF file to tabulate the total number of each SNP class, their genomic distribution, the size of consecutive Diagnostic + or - SNPs, and any SNPs that may be indicative of reciprocal hGC or HE. We developed this script for use cases involving four taxa (*i.e*., two diploid progenitors and two allopolyploid subgenomes) and for cases involving the full seven- taxa patterns (i*.e*., two polyploid subgenome, two diploid progenitors, two diploid outgroups to each subgenome/diploid clade, and an outgroup to the entire genus). This script is available in our Github repository (https://github.com/conJUSTover/GeneConversion). We then explored any difference in the total number of Diagnostic + and - SNPs in R and used ggplot for plotting our results.

### Detecting Potential Regions of Introgression

We used the Dsuite (68) package for inferences of introgression between diploid progenitors, or between polyploid species where we inferred a paraphyletic pattern of hGC. All analyses were done with window sizes of 50 SNPs, and overlapping windows of 10 SNPs.

## ACKNOWLEDGEMENTS

We thank ResearchIT at Iowa State University for computational support. This work was supported by a National Science Foundation Postdoctoral Research Fellowship in Biology (IOS-2209085 to JLC), and National Science Foundation (NSF) awards IOS-1829176 (DBS, JFW, JS, and CEG), IOS-2145811 (JS), and U.S. Department of Agriculture ARS 58-6066-0-066 (DGP, CEG, and JFW).

## Figures and Tables

**Supplementary Figure 1:**
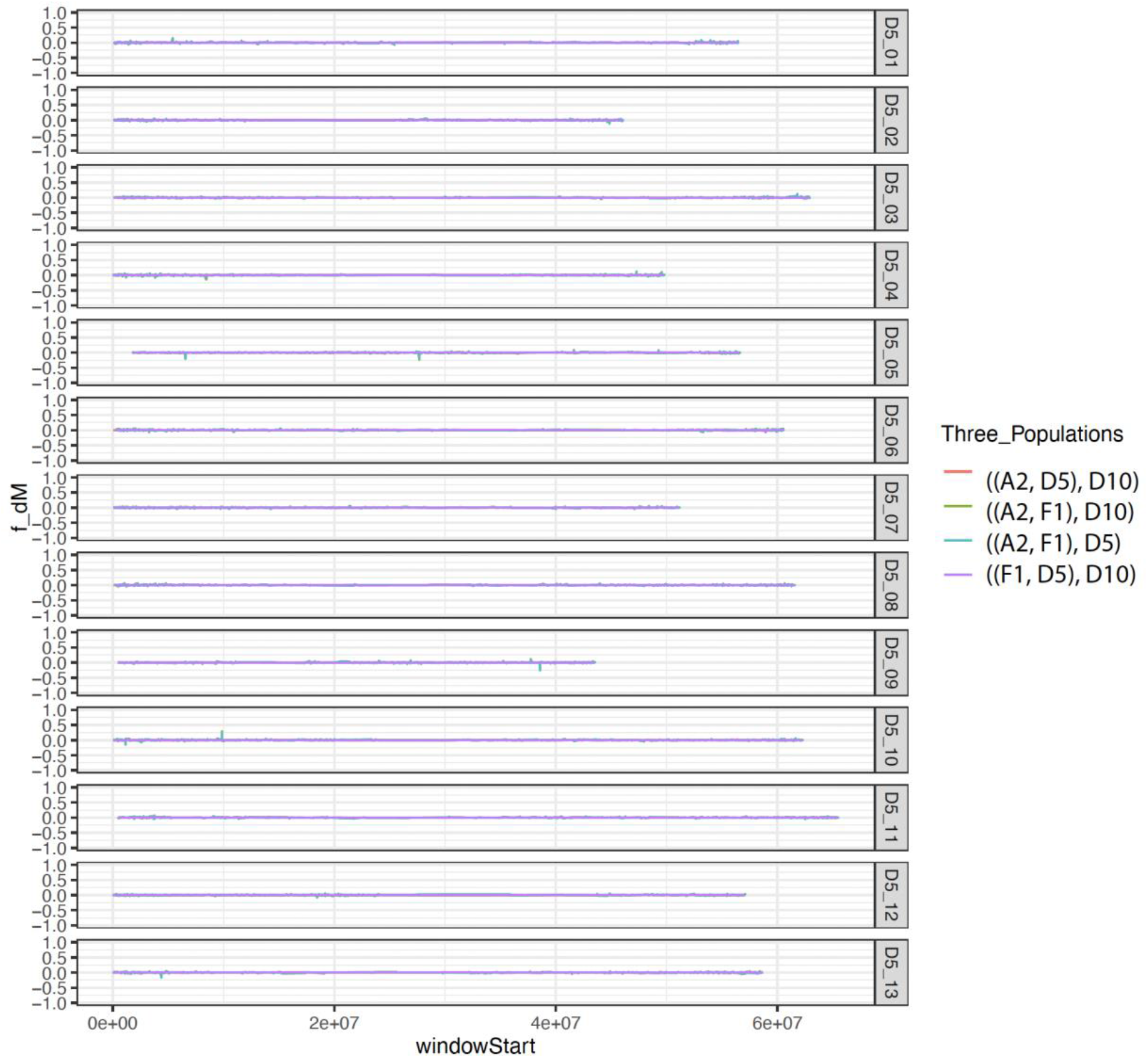
Analyses of introgression between the diploids used in this analysis. Each color indicated a different trio of species, with *Gossypioides kirkii* always serving as the outgroup. Species codes in the legend follow those used in the main text and are as follows: *G. arboreum* (A2), *G. longicalyx* (F1), *G. raimondii* (D5), and *G. turneri* (D10). f_DM values were calculated in Dsuite (68) using window sizes of 50 SNPs and overlapping windows every 10 SNPs.

**Supplementary Figure 2:**
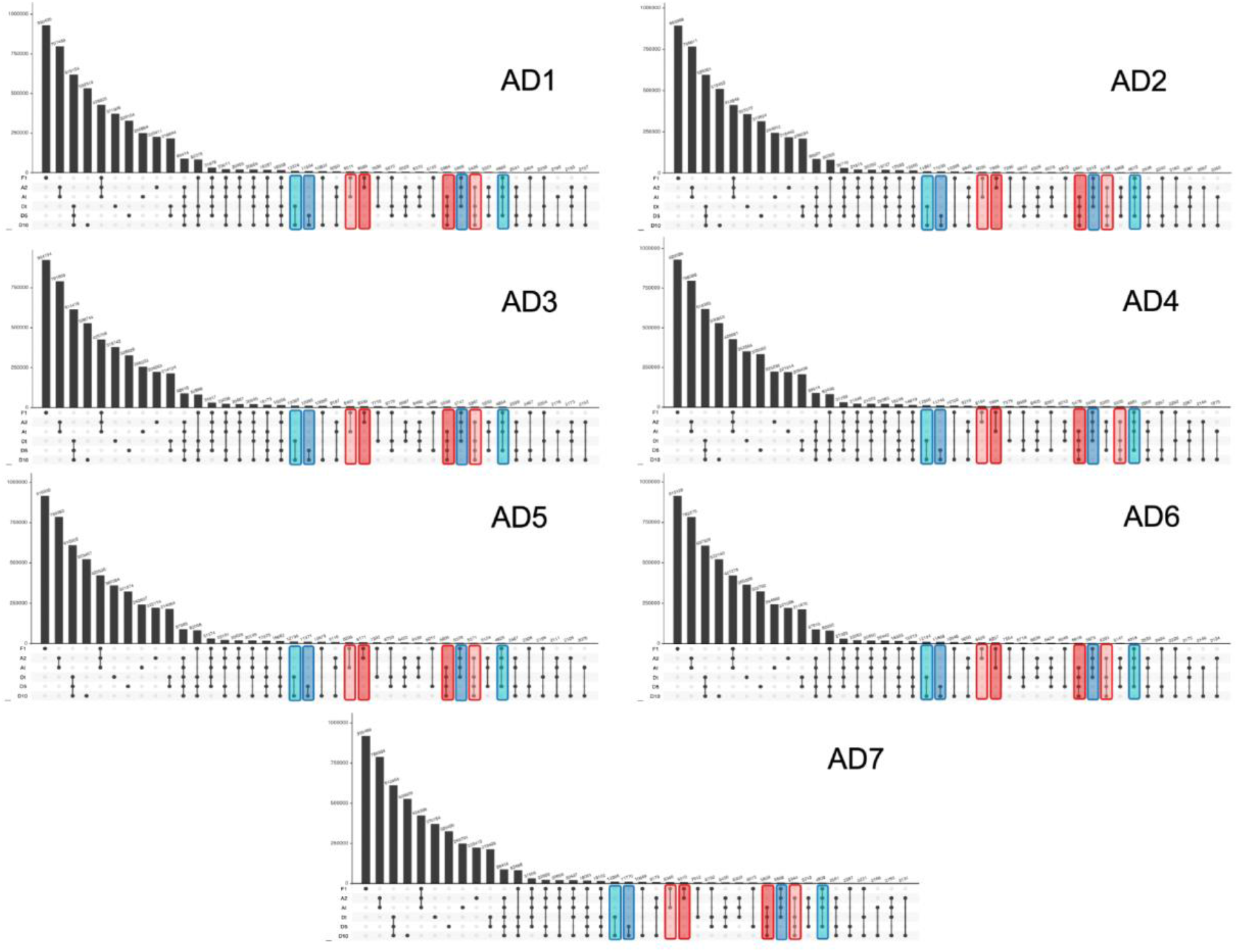
UpSet plots for the top 40 SNP patterns in each of the seven polyploids. Black dots indicate derived alleles, and empty circles indicated ancestral alleles (relative to *Gossypioides kirkii*, which is not shown since all SNPs in this species are inferred to be ancestral). SNPs whos patterns match those that informative as either Diagnostic + (dark blue - Dt overwriting At; dark red - At overwriting Dt) or Diagnostic - (light blue - Dt overwriting At; light red - At overwriting Dt) are highlighted. Polyploid species codes are indicated in the top left of each panel. Species designations in the upset follow the same convention as described in the main text; F1 - G. longicalyx (diploid outgroup to At subgenome); A2 - G. arboreum (diploid progenitor to At subgenome); D5 - G. raimondii (diploid progenitor to Dt subgenome); D10 - G. turneri (diploid outgroup to the Dt subgenome).

**Supplementary Figure 3:**
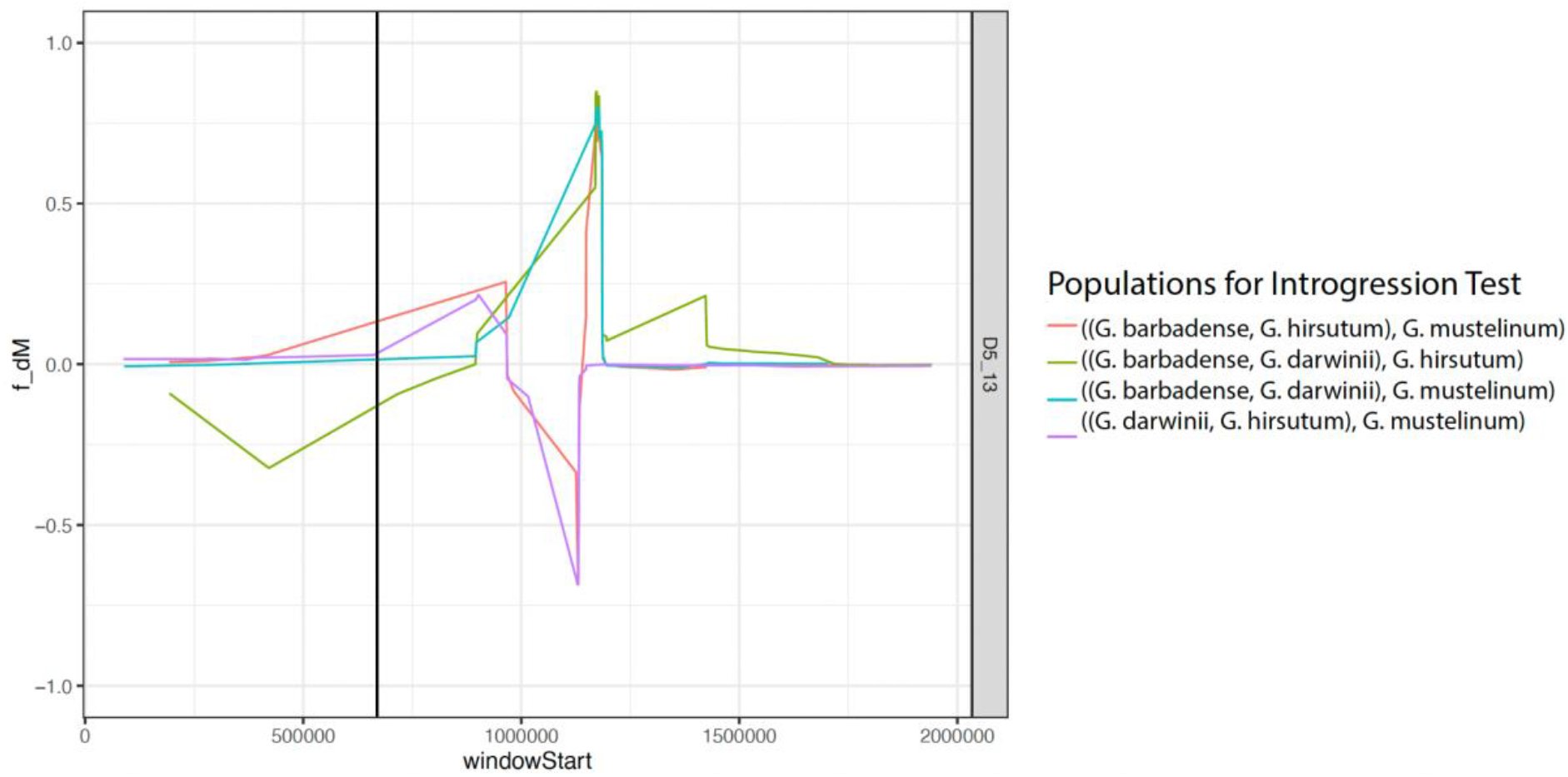
Potential genomic region of introgression from the Dt subgenome of *G. barbadense* into *G. hirsutum* that potentially explains a paraphyletic inference of hGCs in *Gossypium* allopolyploids. Genomic position is indicated on the x axis, with the inferred hGC site indicated by a solid vertical black line. Each color represents a different trio of species shown in Newick format, with *G. raimondii* always serving as the outgroup. f_DM statistic on y axis is the measure of directional selection from (39)in windows of 50 SNPs, with sliding windows every 10 SNPs. All calculations were performed using Dsuite (68).

